# *ImputAccur:* fast and user-friendly calculation of genotype-imputation accuracy-measures

**DOI:** 10.1101/2022.03.07.483268

**Authors:** Kolja A Thormann, Viola Tozzi, Paula Starke, Heike Bickeböller, Marcus Baum, Albert Rosenberger

## Abstract

**Summary:** *ImputAccur* is a software tool for genotype-imputation accuracy-measures. Imputation of untyped markers is a standard approach in genome-wide association studies to close the gap between directly genotyped and other known DNA variants. However high accuracy for imputed genotypes is fundamental. Several accuracy measures have been proposed, but unfortunately, they are implemented on different platforms, which is impractical. With *ImputAccur* the accuracy measures *info, Iam-hiQ* and *r²based* indices can be derived from standard output-files of imputation software. Sample/probe and marker filtering is possible. This allows e.g. accurate marker filtering ahead of data analysis. A Python code is available but also a stand-alone executable file.

**Availability and Implementation:** The source code (Python version 3.9.4), a stand-alone executive file, and example data for *ImputAccur* are freely available at https://gitlab.gwdg.de/kolja.thormann1/imputationquality.git.

**Supplementary Information:** Supplementary information is available at *Bioinformatics* online.

## 1. Introduction

Commercial single nucleotide polymorphism (SNP) microarrays are used to genotype DNA samples for genome-wide association studies (GWAS). Usually, between 300,000 and 4 million variants are genotyped. Imputation methods have been developed to close the gap between genotyped and existing DNA variants (Hickey *et al*., 2013; Das *et al*., 2018; Marchini and Howie, 2010). Most methods estimate a-posteriori genotype probabilities *p*_*g,I,m*_ (one of three possible genotypes *g*) for each untyped SNP/variant/marker *m* and each individual *i* in the sample of interest. The resulting increased variant density improves the genomic coverage and may raise the power to detect associations with a trait (Winkler *et al*., 2014). Quality control of the imputation is essential, e.g. to exclude poorly imputed variants from statistical analysis. Several quality indices have been developed and are routinely applied in studies (Marchini and Howie, 2010; Das *et al*., 2018; Browning and Browning, 2009). These comprise *inter alia* MACH’s *r²*, BEAGLES’s *r²*, IMPUTE2’s *info* or the recently proposed *Iam hiQ*, including a regional classification across markers (Rosenberger *et al*., 2022). Unfortunately, these accuracy measurements are implemented on different platforms. With *ImputAccur* a comfortable use of all these indices is possible. Details and equations of these accuracy indices are summarized in a supplement.

## 2. Format specifications

*ImputAccur* requires the user to provide marker information (leading information) along with the estimated genotype probabilities (dosages) as an input-file, which is a plain text file (or zipped). These are standard files from imputation software. Each row contain information on one marker. The second and third columns should contain [2] a unique marker name and [3] its physical position (e.g. on the chromosome). Probabilities for the genotypes 0, 1 and 2 of each sample/individual can be contained in 3 (summing to 1) or 2 (will amended to one) columns. Missing or inaccurate imputations are indicated by negative values. Hence, the number (no.) of rows in the input-file equals to the no. of genomic markers; the no. of columns equals to the no. of leading columns + 2/3 time the no. of samples/individuals.

An additional input-file for *ImputAccur* should be a parameter-file (params.txt), which is needed to specify the basic settings of the input-files (e.g. no. of leading columns, 2/3 genotype probabilities) and for program control (e.g. name and path of input-file). There is also the option to specify files containing either markers (matching to marker names) or samples/individuals (numbers matching to column order in the input-file) to be excluded from the calculation.

## 3. Launching the application

To invoke the Python code of *ImputAccur* the user may use the following command structure:

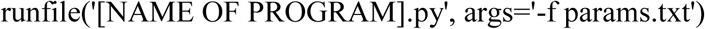

Alternatively, one can run *ImputAccur* as executable file (*ImputAccur*.*exe*) or without the parameter-file (e.g. at a Windows operating system). The program will then ask for entering the parameters interactively.

## 4. Example

Assume your input-file (see example *test1* in the *Supplementary Information*) contains information of 7 SNPs in three leading columns and 3 genotype probabilities each of 5 samples/individuals. Hence, the file has 7 rows and 3+3×5=18 columns. Because the SNP rs00001, rs00003 and rs00004 are quality control markers, these are listed in the file *exclude_SNP*.*txt*. Because individual 1 and 2 are external controls, these are listed in *exclude_PROBE*.*txt*.

This is the input-file (*test1*.*imputed)*:

1 rs00001 913 1 0 0 1 0 0 1 0 0 1 0 0 1 0 0

2 rs00002 402 0.01 0.99 0 0.01 0.99 0 0.01 0.99 0 0.01 0.99 0 0.01 0.99 0

3 rs00003 644 0.333 0.334 0.333 0.333 0.334 0.333 0.333 0.334 0.333 0.333 0.334 0.333 0.333 0.334 0.333

4 rs00004 222 0.25 0.5 0.25 0.25 0.5 0.25 0.25 0.5 0.25 0.25 0.5 0.25 0.25 0.5 0.25

5 rs00005 221 0.47 0.18 0.35 0.89 0.02 0.09 0.03 0.96 0.01 0.94 0 0.06 0.62 0.34 0.04

6 rs00006 955 0.975 0.002 0.023 0.52 0.154 0.326 0.309 0.21 0.481 0.48 0.509 0.011 0.969 0.004 0.027

7 rs00007 518 0.63 0.14 0.23 0.86 0.09 0.05 0.35 0.24 0.41 0.01 0.23 0.76 0.76 0.05 0.19

For this, one need to set the following program parameters in *params*.*txt* or during the execution:

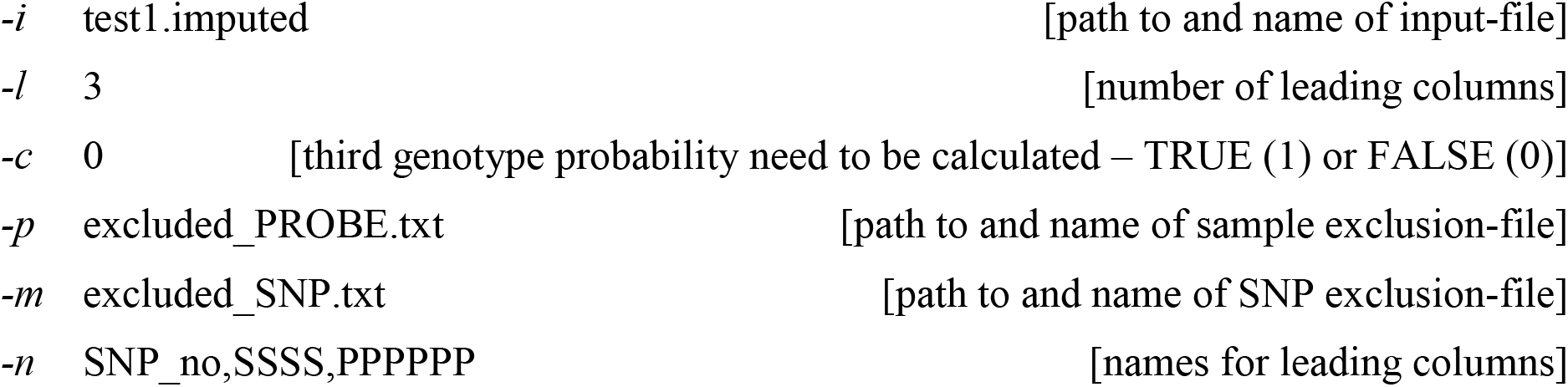

This is the output (*test1*.*accuracy*):

| SNP_no | SNP | position | N | MAF | Iam<sub>chance</sub> | Iam<sub>HWE</sub> | hiQ | accuracy | info | r<sub>2</sub><sub>MACH</sub> | r<sub>2</sub><sub>BEAGLE</sub> |
| --- | --- | --- | --- | --- | --- | --- | --- | --- | --- | --- | --- |
| 5 | rs00005 | 221 | 5 | 26.0% | 0.546 | 0.448 | 0.739 | hot | 0.099 | 0.319 | 0.113 |
| 2 | rs00002 | 402 | 5 | 49.5% | 0.97 | 0.968 | 0.929 | tepid | 0.98 | 0.0 | -9 |
| 7 | rs00007 | 518 | 5 | 40.3% | 0.344 | 0.289 | 0.72 | hot | -0.057 | 0.631 | 0.294 |
| 6 | rs00006 | 955 | 5 | 26.1% | 0.446 | 0.326 | 0.963 | hot | -0.056 | 0.488 | 0.179 |

A regional classification across markers (as outlined by Rosenberger et al. (2022)) is given in the column accuracy.

## 5. Conclusion

*ImputAccur* is an easy-to-use software to determine multiple measures of accuracy for imputed genotypes and independent on the imputation platform used. This allows more flexibility in post-imputation variant filtering. Because it also delivers regional classification, poorly imputed chromosome segments can be identified.

## Supporting information

https://gitlab.gwdg.de/kolja.thormann1/imputationquality/-/blob/development/Supplement_V1.5.pdf

## Code availability

The source code (Python version 3.9.4), a stand-alone executive file and a user-manual of *ImputAccur* are freely available at https://gitlab.gwdg.de/kolja.thormann1/imputationquality.git.

## Data Availability Statement

Example data are available in the same repository as the source code.

