## Supplementary material for "*ImputAccur:* fast and user-friendly calculation of genotype-imputation accuracy-measures": https://gitlab.gwdg.de/kolja.thormann1/imputationquality/-/blob/development/Supplement_V1.5.pdf

*Accuracy measures*

For a di-allelic SNP denote the genotypes *g* by 0, 1 or 2 according to the number of minor alleles A, with minor allele frequency (MAF) $f_A$ in the population. A-posteriori genotype probabilities $p_{g,i,m}$ are given for each untyped SNP/marker *m* and each individual *i*. The *expected* allele dosage of $i^{th}$ probe/individual for the $m^{th}$ SNP/marker is given as $e_{im} = p_{1im} + 2p_{2im}$. We also define $f_{im} = p_{1im} + 4p_{2im}$. To avoid incomputable indices, MAF is estimated as $\widehat{f_A} = \frac{1+\sum_{i=1}^{N} e_{im}}{2+2N}$, with N being the number of samples/individuals, but not less than 1000. The genotypes are in Hardy-Weinberg Equilibrium (HWE) if $[p_0 \quad p_1 \quad p_2] = [f_A^2 \quad 2f_A(1-f_A) \quad (1-f_A)^2]$.

IMPUTE2's *info,* is defined as $$info_m = 1 - \frac{\sum_{i=1}^{N}(f_{im} - e_{im}^2)}{2N\hat{f}_A(1-\hat{f}_A)}.$$

It can be regarded as the proportion of statistical information on MAF in the imputed genotypes, relative to "known" genotypes (Marchini and Howie, 2010 Supplementary Information S3).

MACH $\hat{r}^2$ is defined as $$\hat{r}_m^2 = \left[\frac{\sum_{i=1}^{N} e_{im}^2}{N} - \left(\frac{\sum_{i=1}^{N} e_{im}}{N}\right)^2\right] / 2\hat{f}_A(1-\hat{f}_A).$$

It can be regarded as ratio of the empirically to the expected variance of the allele dosage (under HWE) (Marchini and Howie, 2010 Supplementary Information S3).

BEAGLE $R^2$ is defined as: $$R_m^2 = \frac{\left[\sum_{i=1}^{N} z_{im}e_{im} - \frac{1}{N}\left(\sum_{i=1}^{N} z_{im} \sum_{i=1}^{N} e_{im}\right)\right]^2}{\left[\sum_{i=1}^{N} f_{im} - \frac{1}{N}\left(\sum_{i=1}^{N} e_{im}\right)^2\right]\left[\sum_{i=1}^{N} z_{im}^2 - \frac{1}{N}\left(\sum_{i=1}^{N} z_{im}\right)^2\right]},$$

with $z_{im} \in \{0,1,2\}$ being the most likely imputed genotype *g*. It is the correlation of the best-guess genotype and the allele dosage (Marchini and Howie, 2010 Supplementary Information S3).
*Iam* is a rescaled measure of anti-concentration index $Q_{i,m} = \sum_{g=1}^{3} p_{g,i,m}(1-p_{g,i,m})$, or averages over all marker as $\bar{Q}_m$ (Rosenberger *et al.*, 2022). Because $Q_{i,m}$ can take values between 0 and 2/3 (in the case of equally likely genotypes: $p_{g,i,m}=1/3$),

$\mathrm{Iam}_{\mathrm{chance}}$ is defined as $$Iam_{chance,m} = 1 - \frac{\bar{Q}_m}{2/3}.$$

Considering the genotype probabilities in HWE as natural reference point,

$\mathrm{Iam}_{\mathrm{HWE}}$ is defined as $$Iam_{HWE,m} = 1 - \frac{\bar{Q}_m}{Q_{HWE,m}}.$$

$r^2$-based measures and *info* are directly related to the power of the statistical test of a marker-trait association. Marchini et al. (2010) showed, that *info*, $r^2_{MACH}$ and $r^2_{BEAGLE}$ correlate strongly, but can also exceed 1 or be undefined. In contrast, Rosenberger et al. (2022) showed that *Iam hiQ* and *info* carry different information on imputation accuracy and complement each other as indices.

*Implementation*

*ImputAccur* is a software tool to calculate the genotype-imputation accuracy-measures $Iam_{chance}$, $Iam_{HWE}$, *hiQ, info,* $r^2_{MACH}$ *and* $r^2_{BEAGLE}$, independent of the imputation methods applied. All is needed are dosage files. Additional, whole samples/individuals or SNPs can be in/excluded from the calculation, e.g. to address control samples or markers, double, related or external individuals. Silencing single SNPs per sample/probe from the calculation is also possible. Furthermore, *ImputAccur* classifies markers to be located in a “cold”, “tepid”, “hot” or “very hot” region, the last indicating massive inaccurate imputation, as outlined by Rosenberger et al. (2022).

### *References*


Marchini,J. and Howie,B. (2010) Genotype imputation for genome-wide association studies. *Nat Rev Genet*, **11**, 499–511.

Rosenberger,A. *et al.* (2022) Iam hiQ—a novel pair of accuracy indices for imputed genotypes. *BMC Bioinformatics*, **23**, 50.
